# One-Pot Dual Protein Labelling for Simultaneous Mechanical and Fluorescent Readouts in Optical Tweezers

**DOI:** 10.1101/2024.08.19.608605

**Authors:** Laura-Marie Silbermann, Maximilian Fottner, Nora Migdad, Kathrin Lang, Katarzyna (Kasia) Tych

## Abstract

Optical tweezers are widely used in the study of biological macromolecules but are limited by their one-directional probing capability, potentially missing critical conformational changes. Combining fluorescence microscopy with optical tweezers, employing Förster resonance energy transfer (FRET) pairs, addresses this issue. Moreover, attaching one FRET probe to a tethered protein and the other to a protein in solution allows precise localisation of interaction sites, while probing mechanical properties. When integrating fluorescence microscopy with optical tweezers, orthogonal protein conjugation methods are needed to enable simultaneous, site-specific attachment of fluorophores and DNA handles, commonly used to apply force to molecules of interest. In this study, we utilized commercially available reagents for dual site-specific labelling of the homodimeric heat shock protein 90 (Hsp90) using thiol-maleimide and inverse electron demand Diels–Alder cycloaddition (IEDDAC) bioorthogonal reactions. In a one-pot approach, Hsp90 modified with a cysteine mutation and the non-canonical amino acid cyclopropene-L-lysine (CpK) was labelled with the FRET pair maleimide-Atto550 and maleimide-Atto647N, alongside single-stranded methyltetrazine-modified DNA oligonucleotide. Optical tweezers experiments with this labelled Hsp90 construct revealed structural transitions consistent with previous studies, validating the approach. Fluorescence measurements confirmed the proximity of FRET pairs in the N-terminally closed state of Hsp90 in this experimental setup. This integrative method provides a powerful tool for probing protein conformational dynamics and protein interactions beyond the limitations of traditional optical tweezers.

**Statement:** The developed method combines fluorescence microscopy with optical tweezers, enhancing single-molecule protein studies by overcoming the limitations of one-directional mechanical probing. Utilizing two orthogonal protein conjugation methods for one-pot dual labelling, the heat shock protein 90 was labelled with a FRET pair and single-stranded DNA oligonucleotides. Validated by comparison with published conformational changes, mechanical unfolding signatures, and FRET pair distances, this approach provides a powerful tool to explore single-molecule conformational dynamics and protein interactions.

## 1 INTRODUCTION

A limitation of optical tweezers when applied to the study of biological macromolecules arises from its one-directional probing of structural changes such as unfolding events and conformational changes. This may cause vital information about the studied molecule to go undetected. To overcome this, strategies like engineering multiple force application points within the same molecule to vary the pulling directions (Ziegler et al., 2016) and using a quadrupole-trap (Dame et al., 2006) have been employed. A rapidly growing approach for acquiring comprehensive information about biomolecular systems is the combination of fluorescence microscopy with optical tweezers (Bustamante et al., 2021; Choudhary et al., 2019). By incorporating a Förster resonance energy transfer (FRET) pair into a protein tethered in an optical tweezers setup, it becomes possible to simultaneously detect structural changes that do not occur along the mechanically probed spatial coordinate. Additionally, attaching one FRET probe to the tethered protein and the other to a protein in solution allows for precise localisation of interaction sites, while probing mechanical properties of the studied biological system.

Protein synthesis for simultaneous mechanical and FRET readout in optical tweezers (OT) experiments has previously been achieved using a sophisticated, customized cell-free protein synthesis system to study protein folding within the ribosomal exit tunnel (Wruck et al., 2021). In contrast, our study presents, to our knowledge, the first use of a much more widely adopted *E. coli*-based protein synthesis system to generate protein constructs suitable for simultaneous mechanical and FRET readout in OT experiments.

A crucial consideration for this approach is selecting orthogonal protein conjugation methods for simultaneous, site-specific attachment of fluorophores, and DNA handles commonly used to apply force to a molecule of interest. Decisive factors in choosing two orthogonal protein conjugation strategies include biocompatibility, reagent availability, impact on protein activity, cost, reaction efficiency, and compatibility for a one-pot approach, among others.

Various strategies for attaching DNA handles have been described in recent literature reviews (van der Sleen, 2021; Yang, 2020). When targeting a residue within the protein rather than the N- or C-terminus, conjugation strategies involving single amino acid mutations are preferred, to minimize the risk of affecting protein function. Exploiting the low abundance of cysteines in proteins (< 2%) and their distinctive thiol group for selective reactions (Spicer & Davis, 2014), has been a widely used approach in creating protein-DNA chimeras for optical tweezers experiments. Methods include the simple and fast approach of thiol-maleimide coupling (Jahn et al., 2014). However, the range of functional groups present in natural amino acids is limited, and their abundance poses challenges for achieving site specificity. A solution to this limitation is the use of non-canonical amino acids bearing bioorthogonal handles, whose incorporation into proteins has been extensively optimized in recent years, enabling relatively high yields (Young et al., 2010; Lang & Chin, 2014; Chin, 2017; Scinto et al., 2021).

With these considerations in mind, we chose to use thiol-maleimide chemistry to attach the commercially available FRET pair, maleimide-Atto550 and maleimide-Atto647N (ATTO-TEC, Germany). Additionally, we employed the irreversible and relatively rapid inverse electron demand Diels-Alder cycloaddition (IEDDAC) (Mattheisen et al., 2023; Pagel, 2019) for the reaction between the commercially available single-stranded methyltetrazine-modified DNA oligonucleotide (e.g., from Biomers, Germany), more specifically a DNA oligonucleotide functionalized with a 3’-aminolink-C6 and 2-(4-(6-Methyl-1,2,4,5-tetrazin-3-yl)phenyl)acetic acid, and the non-canonical amino acid N-(((2-methylcycloprop-2-en-1-yl)methoxy)carbonyl)-L-lysine (cyclopropene-L-lysine, CpK) (e.g., from SiChem GmbH, Germany) (Elliott et al., 2014). This non-canonical amino acid was incorporated into the Hsp90 protein from *Saccharomyces cerevisiae* (Hsp82). Combining mutually orthogonal IEDDAC and thiol-maleimide labelling in a one-pot setup has already been successfully demonstrated using a small molecule substrate (Knall et al., 2016). Although Hsp90 remains enigmatic in many ways (Silbermann, Vermeer et al., arXiv:2308.16629), it has already been extensively studied in optical tweezers experiments (Jahn et al., 2014, 2016, 2018; Tych et al., 2018) making it a suitable model protein for developing an optical tweezers method involving a tethered, fluorescently labelled protein.

## 2 RESULTS

### Generating Hsp90 protein with two site-specific chemical attachment sites

To allow for simultaneous, site-specific labelling of Hsp90 with commercially available fluorescent dyes and short single-stranded DNA oligonucleotides, we inserted two chemical attachment sites. This was achieved by site-directed mutagenesis. The amino acid sequence of the construct is provided in the Supplementary Information (SI). One attachment site was introduced by creating a cysteine mutation in the N-terminal domain (NTD) at amino acid position 61, and the other by introducing an amber stop codon at amino acid position 452 in the middle domain (MD) for incorporation of the non-canonical amino acid CpK via genetic code expansion (Figure 1A, B). Both the cysteine at position 61 and CpK at 452 are solvent exposed. Another cysteine mutation was inserted into an artificial C-terminal coiled-coil motif at position 6. The C-terminal coiled-coil motif, which has been previously reported and utilized, stabilizes the dimeric form of Hsp90. The cysteines in the coiled-coil motif are closely spaced, promoting disulfide bond formation, covalently linking the monomers and facilitating re-dimerization during repeated pulling and relaxation cycles in optical tweezers experiments. Cysteines at position 61 are further apart and less likely to be involved in disulfide bond formation (Jahn et al., 2016). Positions 61 in dimeric Hsp90 are suitable for labelling with a FRET pair, as previous single-molecule FRET experiments revealed that these positions are in close enough proximity in the N-terminally closed state (87.8 Å) but not in the N-terminally open state ensemble for FRET to occur. Inter-domain fluctuations of up to 25 Å have been reported for the ensemble of open state conformations in the absence of an applied force (Hellenkamp, et al. 2017).

**Figure 1:**
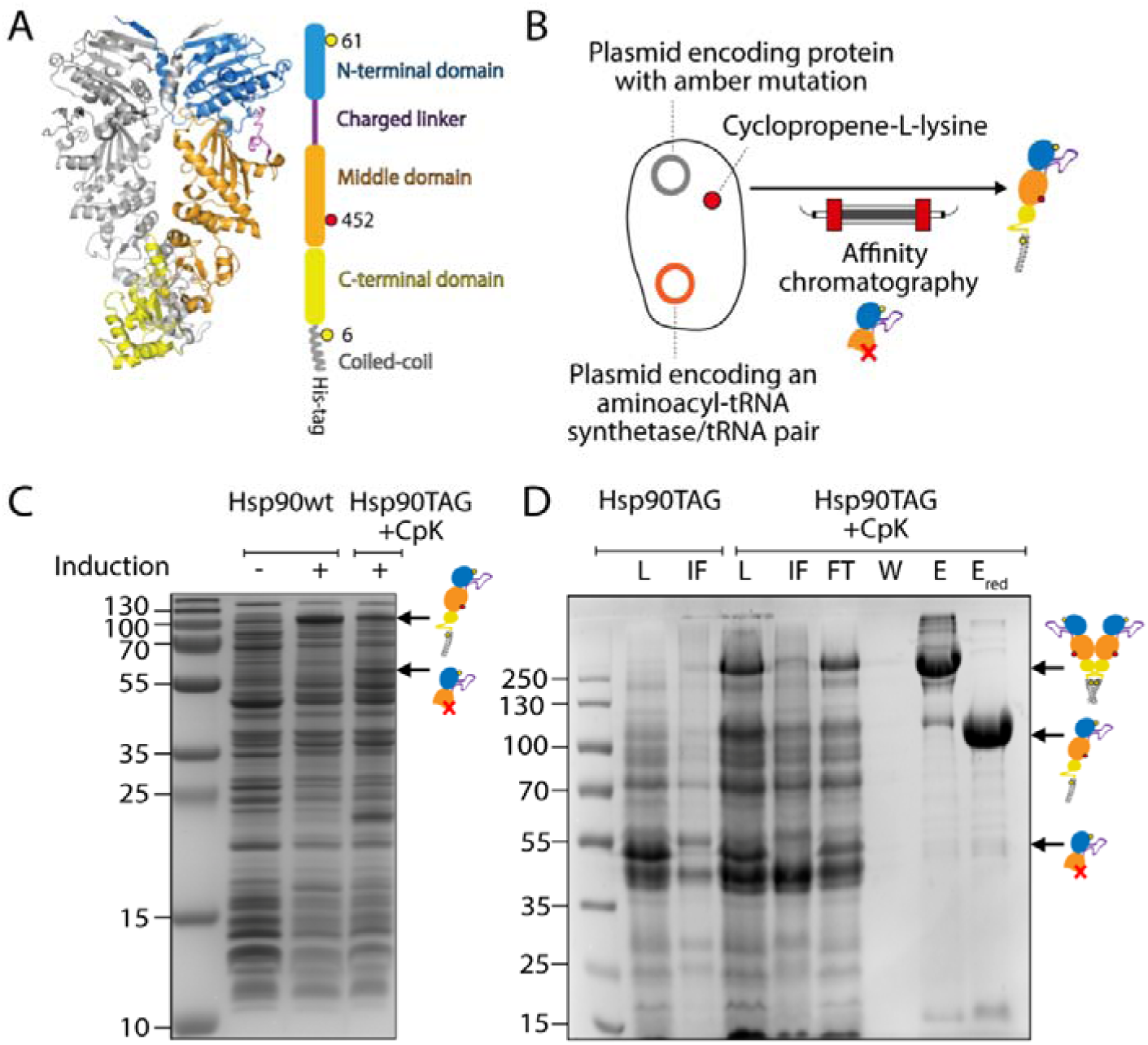
Producing Hsp90 protein featuring two site-specific chemical attachment sites for labelling. (A) The crystal structure (PDB accession code: 2CG9) is shown with one monomer coloured by its different structural elements: the NTD (blue), charged linker – not fully resolved in the crystal structure – (purple), MD (orange), C-terminal domain (CTD, yellow) and coiled-coil (grey). Introduced modifications are cysteines at position 61 (D61C) and 6 (A6C) (yellow circles) as well as an amber stop codon at position 452 (D452TAG, red circle). (B) Expression was performed using *E. coli* K12 cells harbouring two plasmids: one with the Hsp90 construct, and the other plasmid with an aminoacyl-tRNA synthetase (aaRS)/tRNA pair that directs the incorporation of CpK in response to the amber codon. Truncated protein resulting from incomplete amber suppression was removed by affinity chromatography using a C-terminal affinity tag. (C) SDS-PAGE gel of cell lysate supernatant from Hsp90wt and Hsp90-D452TAG under various expression conditions (D) SDS-PAGE gel of samples from Ni-IMAC purification include: load (L), flow through (FT), insoluble fraction (IF), wash (W), eluate (E) and eluate with reducing agent (E_red_).

CpK, added to the culture medium of *E. coli* K12 cells co-transformed with two plasmids - one carrying the modified Hsp90 and the other carrying an orthogonal aminoacyl-tRNA synthetase (aaRS)/tRNA pair for CpK encoding - was successfully incorporated at position 452 via amber (TAG) stop codon suppression. Amber suppression of Hsp90-D452TAG with CpK showed good yields, when compared to Hsp90wt, and only little amounts of truncated protein (52.3 kDa) (Figure 1 C). Truncated Hsp90 was successfully removed by Ni-IMAC affinity chromatography (lanes E and E_red_ in Figure 1D) utilizing the C-terminal His-tag on full-length Hsp90. Additional SEC purification enabled the isolation of protein of high purity. From a 1 L culture, 1.8 mg of purified protein was obtained, which was subsequently used for labelling with FRET pair dyes and short single-stranded DNA oligonucleotides.

### Labelling with DNA oligonucleotides and a FRET pair using a one-pot approach

Labelling was performed using an efficient one-pot approach (Figure 2A) that combines two mutually orthogonal chemistries. One of the chemistries involves the IEDDAC between the non-canonical amino acid CpK in the Hsp90 MD and single-stranded methyltetrazine-modified DNA oligonucleotide. The other uses a thiol-maleimide reaction to label the cysteine at position 61 in the NTD of dimeric Hsp90 with the FRET pair Atto-550 and Atto-647N. The thiol-maleimide reaction requires cysteine reduction prior to adding labelling reagents (Figure 2B). The removal of the reducing agent (here TCEP) before dimerization and labelling was initially attempted using a MiniTrap G-25 column, known for its higher step yield compared to the larger SEC columns typically used for protein purification. However, using a MiniTrap G-25 column negatively affected the degree of covalent dimerization, leading to a switch to a Superdex 200 column. After labelling, free oligonucleotides, dyes, and aggregated proteins were removed by SEC.

**Figure 2:**
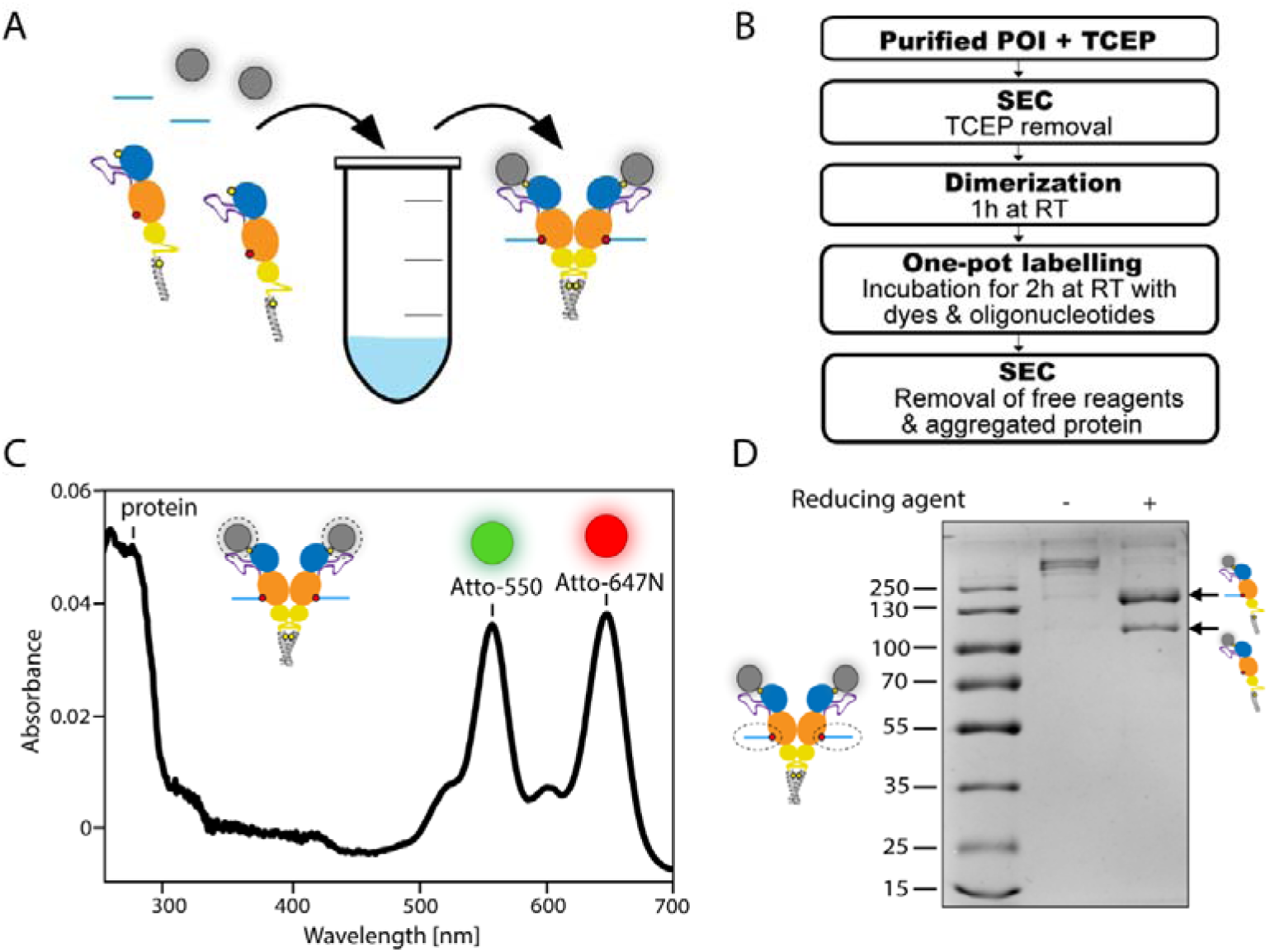
Labelling with DNA oligonucleotides and a FRET pair using a one-pot approach. (A) One-pot functionalization with single-stranded methyltetrazine-modified DNA oligonucleotide via reaction with the non-canonical amino acid CpK, and attachment of the maleimide-Atto550 and maleimide-Atto647N FRET pair to cysteine. (B) Functionalization of the protein of interest (POI) and purification scheme. (C) Degree of labelling with fluorescent dyes determined spectrophotometrically. (D) Degree of labelling with DNA oligonucleotide determined by SDS-PAGE analysis and protein band quantification using ImageJ.

Subsequently, the degree of labelling with dyes (DOL_dye_) was determined spectrophotometrically (Figure 2C) as described in the Methods section. The DOL_dye_ was found to be 0.5 for Atto-550 and 0.4 for Atto-647N, indicating that 90% of purified Hsp90 monomers were labelled with dyes in an almost equal dye ratio. This high labelling efficiency, resulted in 40% of the dimeric Hsp90 containing a FRET pair.

Labelling efficiency with single-stranded methyltetrazine-modified DNA oligonucleotide (DOL_oligo_) was determined to be 69% using an SDS-PAGE-based approach (Figure 2D).

### Mechanical readout in optical tweezers experiments

Dissociation and unfolding of the Hsp90 dimer under force applied at amino acid position 452 in the MD has been previously characterized using an optical tweezers setup (Tych et al., 2018). This study observed reproducible changes in contour length, attributed to the dissociation and unfolding of the CTD and the partial unfolding of the middle domain (MD_part_, amino acids 453–527), by comparing these changes to the Hsp90 crystal structure (Tych et al., 2018). In our study, optical tweezers experiments were conducted with the same Hsp90 construct, using the non-canonical amino acid CpK at position 452 instead of cysteine, and with position 61 of dimeric Hsp90 labelled with a FRET pair (Figure 3A).

**Figure 3:**
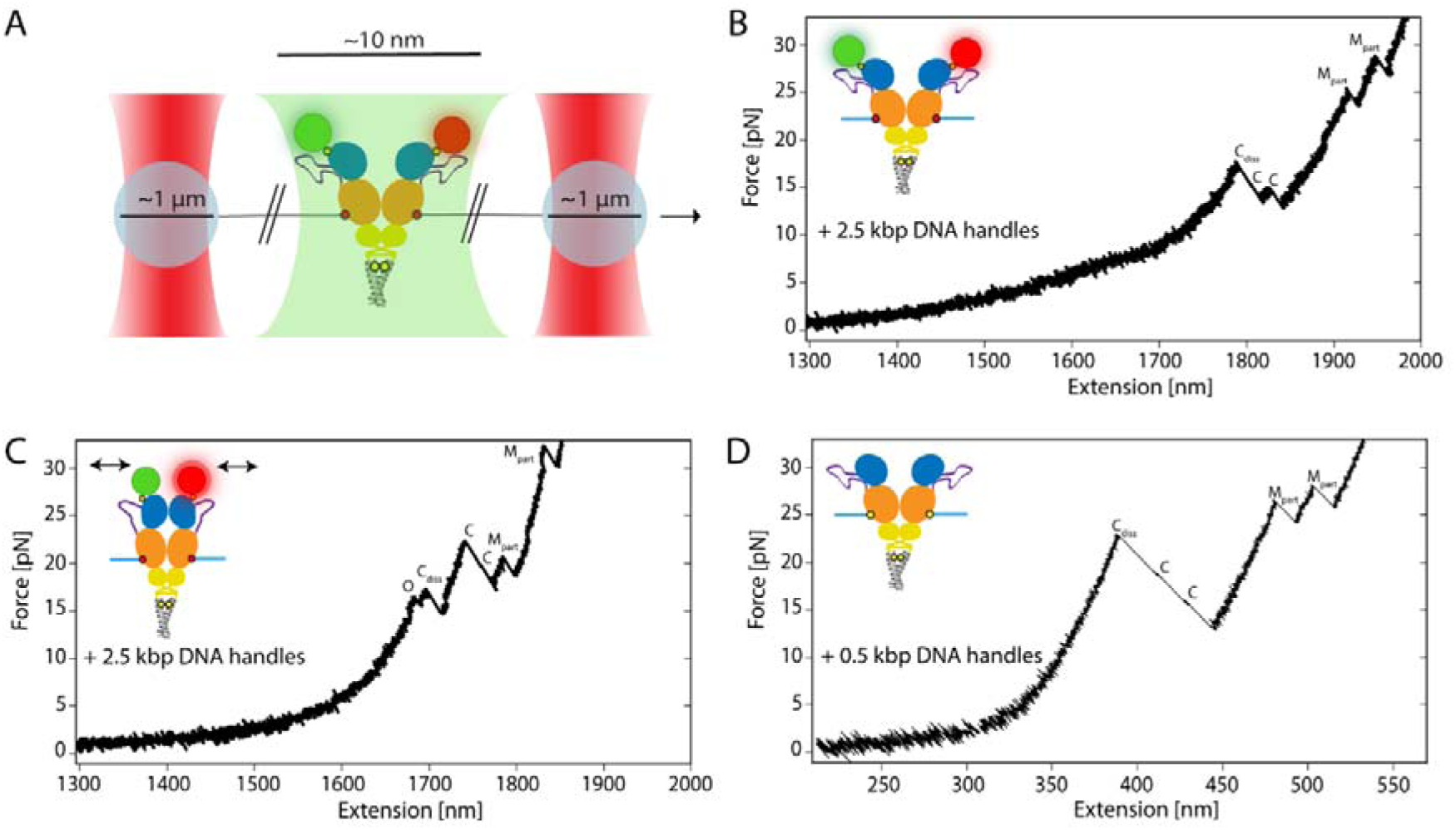
Force readout of fluorescently labelled Hsp90 in optical tweezers experiments. (A) Schematic illustration of the experimental setup: Hsp90 labelled with a FRET pair is tethered between two trapped polystyrene beads using 2.5 kbp-long DNA handles. Hsp90 domains are displayed in different colours: NTD (blue), linker (purple), MD (orange), CTD (yellow), and the artificial coiled-coil motif (grey). Introduced non-canonical amino acid CpK (red circle) and cysteines (yellow circle) with attached fluorescent dyes Atto-550 (green circle) and Atto-647N (red circle). Excitation laser light is depicted in green. (B) Example unfolding trace of fluorescently labelled Hsp90 in a constant velocity (500 nm/s) optical tweezers experiment. The trace shows the initial dissociation of the Hsp90 CTDs (C_diss_), followed by the unfolding of both Hsp90 CTDs (C), and finally the unfolding of both Hsp90 MDs (M_part_). (C) Similar to (B), but using fluorescently labelled Hsp90 in presence of AMP-PNP, known to promote the formation of the N-terminally closed state of Hsp90. The trace shows an additional length change resulting from the transition from the N-terminally closed to the open conformation (O), followed by the dissociation of the Hsp90 CTDs (C_diss_), unfolding of both Hsp90 CTDs (C), and finally the unfolding of both Hsp90 MDs (M_part_). (D) Similar to (B), but using non-fluorescently labelled Hsp90 and 0.5 kbp-long DNA handles.

Measurements were conducted, both in the absence and presence of the nucleotide AMP-PNP, a known inducer of the closed state of the N-terminal domains of the dimeric Hsp90 (Ratzke et al., 2014), resulting in 147 unfolding traces obtained in the absence of AMP-PNP and 26 traces in its presence, examples of which are shown in Figures 3B and 3C. Observed events, along with the corresponding contour length gains, as well as dissociation and unfolding forces of the construct labelled with fluorescent dyes, are comparable to already published data of the Hsp90 construct without attached dyes (Tych et al., 2018), except for the force at which dissociation of the C-terminal domains occurs (Figure 3B-D and Table 1). This force is reduced for the dye-labelled construct in this study.

**Table 1:**
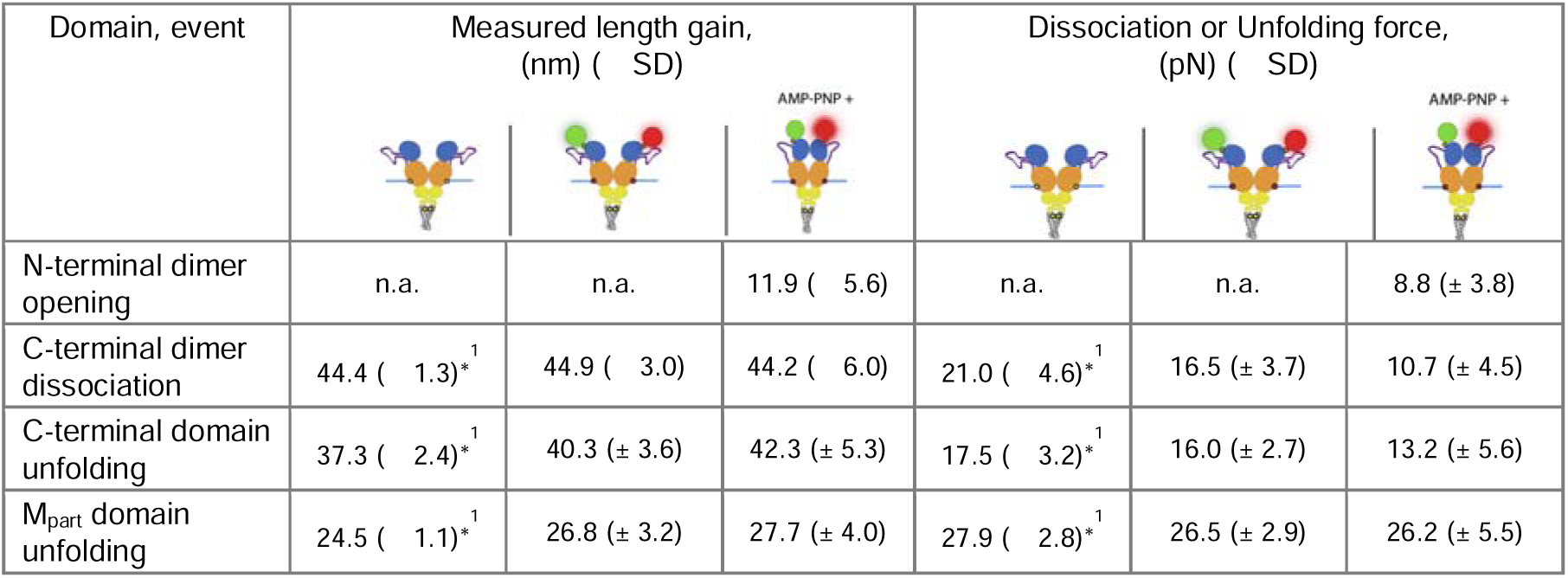
Table of measured length gains and dissociation or unfolding forces of the fluorescently labelled middle-domain attachment construct, compared to previously published data*^1^ (Tych et al., 2018) for the non-fluorescently labelled construct.

| Domain, event | Measured length gain,<br>(nm) ( SD) |  |  | Dissociation or Unfolding force,<br>(pN) ( SD) |  |  |
| --- | --- | --- | --- | --- | --- | --- |
| N-terminal dimer opening | n.a. | n.a. | 11.9 ( $\pm 5.6$ ) | n.a. | n.a. | 8.8 ($\pm 3.8$) |
| C-terminal dimer dissociation | 44.4 ( $\pm 1.3$ )<sup>*1</sup> | 44.9 ( $\pm 3.0$ ) | 44.2 ( $\pm 6.0$ ) | 21.0 ( $\pm 4.6$ )<sup>*1</sup> | 16.5 ($\pm 3.7$) | 10.7 ($\pm 4.5$) |
| C-terminal domain unfolding | 37.3 ( $\pm 2.4$ )<sup>*1</sup> | 40.3 ($\pm 3.6$) | 42.3 ($\pm 5.3$) | 17.5 ( $\pm 3.2$ )<sup>*1</sup> | 16.0 ($\pm 2.7$) | 13.2 ($\pm 5.6$) |
| MD<sub>part</sub> domain unfolding | 24.5 ( $\pm 1.1$ )<sup>*1</sup> | 26.8 ($\pm 3.2$) | 27.7 ($\pm 4.0$) | 27.9 ( $\pm 2.8$ )<sup>*1</sup> | 26.5 ($\pm 2.9$) | 26.2 ($\pm 5.5$) |

This difference is likely due to the use of approximately five times longer DNA handles (2.5 kbp) giving the domains more time to dissociate before the force increases to higher values (Oberbarnscheidt et al., 2009). Longer handles were used to increase the distance between the dyes and the high-intensity trapping lasers to minimize photobleaching. In conclusion, the attached dyes do not appear to affect Hsp90’s conformation, probed domain stabilities, or refolding capacity. In the presence of AMP-PNP, an additional observation was made: a reproducible contour length gain of 11.9 nm (± 5.6 nm) at a low force of 8.8 pN (± 3.8 pN) across various traces (event "O" in Figure 3C). This event has also been observed for unlabelled Hsp90 (Figure S1) and consistently occurred before the dissociation and unfolding of the Hsp90 CTDs and the unfolding of the MD_part_ domains, likely due to the opening of the N-terminally closed dimer. In single-molecule FRET experiments (Hellenkamp et al., 2017), the distance between residues at position 452 was found to be 6.1 nm in the closed state and 7.9 nm in the open state of Hsp90, demonstrating an increase of 1.8 nm (7.9 nm - 6.1 nm) with N-terminal dimer opening.

Upon applying force, an unstructured region connecting the Hsp90MD and CTD, as well as the loop region around the force application point, contributes an additional extension of 5.5 nm per Hsp90 monomer. Consequently, the total expected increase in length from the closed state of Hsp90 to the open state, with the C-terminal dimerization interface intact, is estimated at approximately 12.8 nm (5.5 nm × 2 + 1.8 nm). This approximation matches well with the observed contour length gain of 11.9 nm (± 5.6 nm).

### Fluorescence readout in optical tweezers experiments

To enable the detection of fluorescent dyes attached to Hsp90 with the hybrid C-Trap system, it is crucial to accurately align the confocal focus with the optical trap’s focal plane. This procedure is detailed in the Supplementary Information (Figure S2). To monitor fluorescence along a selected scan line over time during repeated stretching and relaxation cycles, fluorescence data were acquired through kymograph recordings (Figure 4A). In the N-terminally open state of Hsp90, the two fluorescent dyes are too distant for FRET to occur. Measurements were therefore also performed in the presence of AMP-PNP, which strongly shifts Hsp90 to an N-terminally closed state. This conformation brings the fluorescent dyes into close proximity, allowing FRET to occur (Figure 4B). Upon the application of force to Hsp90, a decrease in red fluorescence and an increase in green fluorescence were observed. This indicates a transition from the N-terminally closed conformation, with the FRET pair in close proximity, to the N-terminally open conformation with the FRET pair too far apart for FRET to occur. For clarity, Figure 4 shows only the scanned area of the tethered protein. Kymograph recordings, including the scanned area of the beads, are presented in Figure S3.

**Figure 4:**
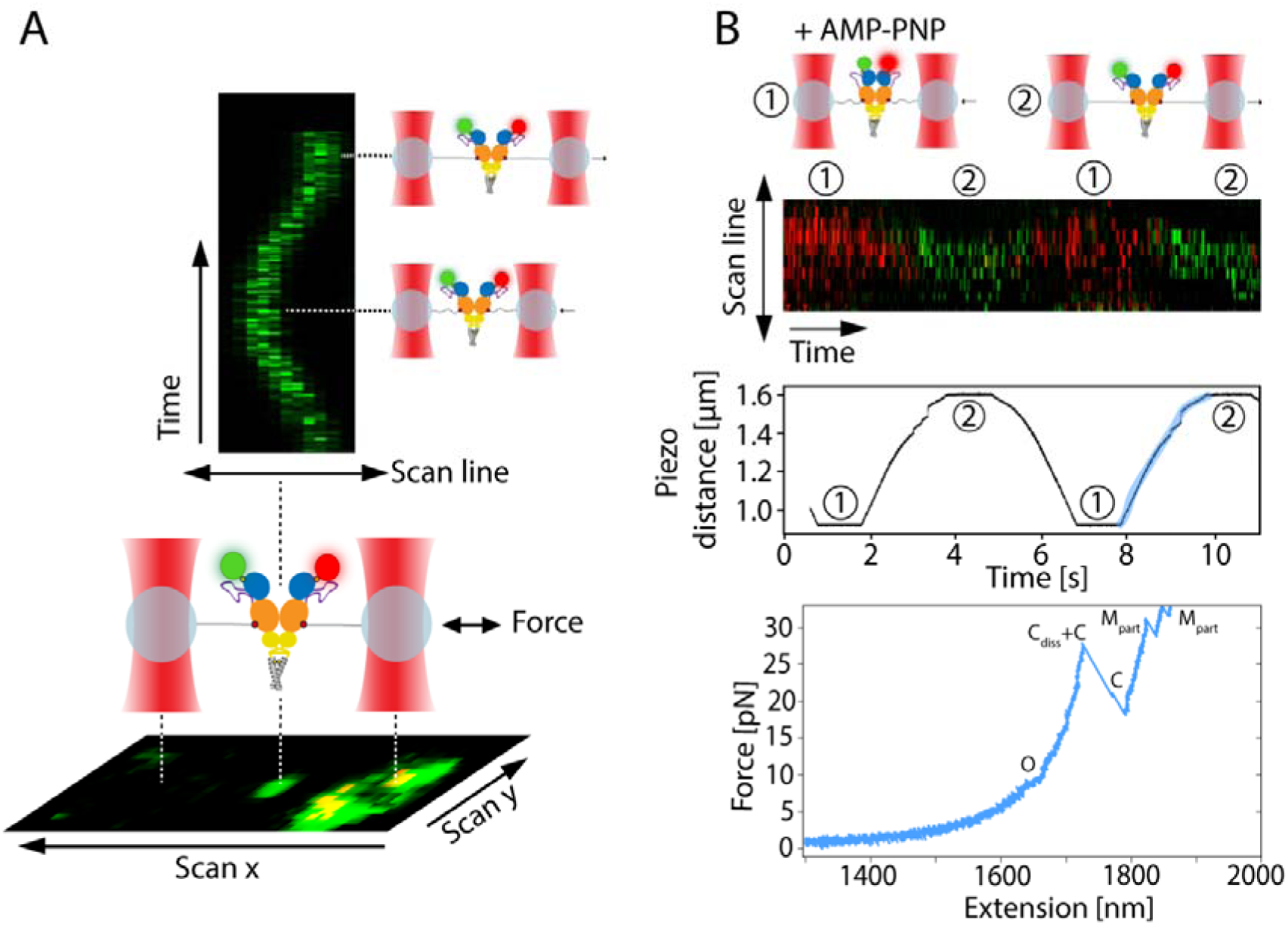
Kymograph recordings. (A) Illustration of an image scan (bottom) and example kymograph recording (top) depicting Hsp90 labelled with the FRET pair Atto-550 (green circle) and Atto-647N (red circle) during repeated stretching and relaxation cycles in the absence of AMP-PNP. (B) Example kymograph recorded in the presence of AMP-PNP (top) with the piezo distance versus time plot from the same recording below. The stretching cycle region of the piezo distance graph, corresponding to the displayed unfolding trace (lowest panel), is highlighted in blue. The force-extension readout indicates the transition from the N-terminally closed to open conformation (O), followed by the dissociation of the Hsp90 CTDs (C_diss_), unfolding of both CTDs (C), and finally the partial unfolding of both MDs (M_part_). The instrument used had an offset between the green and red channels. To correct this in the displayed kymographs, a custom Python script was used to determine the pixel shift between the two channels. The shift was then applied using Photoshop, which was also used to adjust the brightness for better visualization.

## 3 DISCUSSION

Optical tweezers have revolutionized the study of biological macromolecules by enabling precise force manipulation at the single-molecule level. However, their traditional one-directional probing capability limits the comprehensive characterization of conformational dynamics. These limitations can be overcome by integrating fluorescence microscopy with optical tweezers. In this study, we develop a method to enable the observation of conformational changes and mechanical unfolding of a tethered protein labelled with a FRET pair using a system that has already been well characterized in OT experiments – Hsp90 (Jahn et al., 2014, 2016, 2018; Tych et al., 2018).

Central to the success of this integrative approach is the use of selective and efficient orthogonal protein conjugation methods that preserve protein function. In this study, genetic code expansion, coupled with a cysteine mutation, was employed to create specific chemical attachment sites. An optimized system for incorporating the non-canonical amino acid CpK resulted in a yield of 1.8 mg of purified protein from a 1 L culture. This amount of protein is sufficient to perform a large number of optical tweezers experiments. Labelling of Hsp90 was achieved in a one-pot approach using orthogonal chemistries, resulting in a high DOL with both fluorescent dyes, consistent with previous reports (Gebhardt et al., 2021), and single-stranded DNA oligonucleotides. The Hsp90 construct labelled in this manner exhibited structural transitions and dissociation/unfolding forces consistent with those observed in a previous optical tweezers study (Tych et al., 2018).

The attached FRET pair confirmed the proximity of the labelled residues in the N-terminally closed state of Hsp90, consistent with existing single-molecule FRET data (Hellenkamp et al., 2017). This validation not only confirms the experimental approach but also highlights its utility in studying biomolecular conformational transitions. Expanding this approach to other protein systems that exhibit internal correlated motions along multiple coordinates, such as membrane proteins (van der Sleen et al., 2024; Yang et al., 2018) and kinases (Xie et al., 2020; Gough & Kalodimos, 2024), will allow to provide deeper insights into their complex conformational dynamics. Additionally, this approach can be employed to precisely locate protein interaction sites in an optical tweezers setup by attaching one FRET probe to the tethered protein and the other to a protein in solution, allowing for the simultaneous detection of changes in mechanical properties, such as stretching-induced interactions of proteins involved in mechanotransduction (Del Rio et al., 2009) and the modulation of protein structural stability upon binding an interaction partner (Dahal et al., 2020; Mondol et al., 2023).

## 5 CONCLUSIONS

In conclusion, the integration of fluorescence microscopy with optical tweezers, supported by orthogonal protein labelling strategies, provides a powerful tool for studying conformational dynamics and protein interactions at the single-molecule level. The use of an *E. coli*-based protein synthesis system to generate protein constructs, combined with a one-pot dual labelling approach using commercially available reagents, ensures both efficiency and ease of implementation. By overcoming the limitations of traditional force spectroscopy, this approach opens new avenues for exploring biomolecular systems that exhibit conformational dynamics along multiple coordinates. Additionally, it allows for the precise localisation of protein interaction sites while simultaneously providing valuable insights into the mechanical properties of the studied system.

## 6 MATERIALS AND METHODS

### Expression

Chemically competent *E. coli* K12 underwent co-transformation with two plasmids: a pBAD plasmid carrying Hsp90 with a D61C mutation and an amber stop codon at D452, and a pEVOL plasmid carrying *Methanosarcina barkeri (Mb)* wt pyrrolysyl-tRNA synthetase and the corresponding *Mb* tRNA.

Following recovery in 1 mL of SOC medium for an hour at 37 °C, the cells were cultured overnight in 50 mL of 2×YT medium supplemented with chloramphenicol (25 μg/mL) and ampicillin (50 μg/mL) at 37 °C and 180 rpm. This overnight culture was then diluted to an OD_600_ of 0.05 in 200 mL fresh 2×YT medium with the same supplements and incubated at 37 °C with agitation at 180 rpm until reaching an OD_600_ of 0.6. Subsequently, 1 mM of CpK (SiChem GmbH, Germany) and 0.02% (w/v) of arabinose were added, inducing protein expression for 16 h at 20 °C. Finally, the cells were harvested via centrifugation at 8,000 g for 20 min at 4 °C.

### Protein preparation

Cells were lysed by sonication in a buffer containing 40 mM HEPES (pH 7.5), 150 mM NaCl, 10 mM imidazole, 1 mM MgCl_2_ with an EDTA-free protease inhibitor cocktail added. Protein preparations underwent a multi-step protocol: first, Ni-IMAC (Ni-NTA Agarose, Qiagen, Germany) performed at 4°C to minimize protease activity, followed by size exclusion chromatography (SEC) (Superdex 200, Cytiva, Germany). Fractions of highest purity were combined and incubated with 10 mM TCEP for 30 min at room temperature (21°C) in the dark to reduce cysteines. TCEP removal was achieved via SEC (Superdex 200, Cytiva, Germany), followed by a 1 h room temperature (21°C) incubation for dimerization, covalently linking formed homodimers through a disulfide bond in the artificial coiled-coil motif. Next, protein in PBS (pH 6.7) buffer was labelled in a one-pot approach with a 3-fold molar excess of an equimolar mix of maleimide-Atto550 and maleimide-Atto647N FRET pair (ATTO-TEC, Germany) along with 3-fold molar excess of single-stranded methyltetrazine-modified DNA oligonucleotide (Biomers, Germany) for 2 h at room temperature (21°C) in the dark. The maleimide group, reactive toward solvent-exposed cysteines, enabled dye attachment at position 61. DNA oligonucleotides were attached at position 452 via IEDDAC of methyltetrazine with the non-canonical amino acid CpK. Finally, SEC was used to remove free oligos, dyes, and aggregated protein, and to exchange the buffer to 40 mM HEPES (pH 7.5), 150 mM KCl, 10% glycerol.

### Determination of the average degree of labelling

DOL_dye_, which refers to the average number of dye molecules attached to each protein molecule, was determined for the conjugate solution obtained after the final SEC step. This was accomplished using absorption spectroscopy and applying the Lambert-Beer law using the theoretical extinction coefficient of the monomeric Hsp90 construct (ε_*prot*_ =75415 M^-1^ cm^-1^). Absorbance measurements were taken at the dyes’ absorption maxima (A_max_) as well as at the protein’s absorption wavelength, 280 nm (A_prot_). Since all dyes exhibit absorbance at 280 nm, the measured absorbance at this wavelength (A_280_) was corrected to account for the dyes’ contribution. This correction followed the method outlined by the dye supplier ATTO-TEC, utilizing the provided correction factors (*CF*_280_) and dye extinction coefficients (*ε*_*max*_):

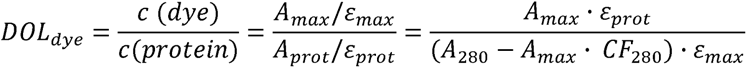

DOL_oligo_ was determined by quantifying the bands of oligonucleotide-labelled and non-oligonucleotide-labelled protein separated on SDS-PAGE gels using ImageJ.

### Attachment of DNA handles

The DNA oligonucleotides bind to the sticky ends of 2.5 kbp (∼1.7 µm) DNA handles, functionalized with digoxigenin or biotin. These handles were synthesized in-house using Lambda DNA as a template (New England Biolabs, USA) and custom-designed primers (Metabion, Germany). This was done similarly to synthesis of shorter 0.5 kbp handles used for the non-fluorescently labelled construct, which is described in Tych & Rief (2022). DNA handles were incubated with oligo-protein in the presence of 2 mM MgCl_2_ for 1 h at room temperature (21°C) and 150 rpm on a plate shaker.

### DNA sequences

DNA oligonucleotide modified with a methyltetrazine group

5’-G GCA GGG CTG ACG TTC AAC CAG ACC AGC GAG TCG-methyltetrazine-3’

Custom-designed primers used to make DNA handles

5’-Biotin-GTAC(Biotin-dT)GGA(Biotin-dT)GCACTG GAG AAG-3’

5’-Digoxigenin-GTAC(Dig-dT)GGA(Dig-dT)GCACTG GAG AAG-3’

5’-CGACTCGCTGGTCTGGTTGAACGTCAGCCCTGCC(abasic site)CCTGCCCGGCTCTGGACAGG-3’

### Optical tweezers experiments

Optical tweezers experiments using a C-trap instrument (Lumicks) that combines a confocal microscope with optical tweezers were performed as described before (Mondol et al., 2023), but using 0.75 µm antidigoxigenin and 1.36 µm streptavidin-coated polystyrene beads and conducting the measurements at a constant velocity of 500 nm/s instead. In addition to the used oxygen scavenger system – comprising 1700 U/mL glucose catalase, 27 U/mL glucose oxidase, and 0.66% glucose to minimize photodamage caused by oxygen free radicals – 1 mM Trolox was included in the measurement buffer as an antiblinking and antibleaching reagent.

To align the confocal focus (determined by the objective position) with the trap focal plane, where the beads are trapped with the tethered sample, SYBR Safe stain (Invitrogen, USA) was employed at a final concentration of 250 × (achieved by diluting the 10,000 × stock solution 40 times).

Measurements with fluorescently-labelled Hsp90 construct were additionally performed in the presence of 2 mM AMP-PNP, following incubation with the sample for 30 min at room temperature (21°C). Fluorescence data were collected using the hybrid Lumicks instrument’s confocal system, which is equipped with three excitation lasers (488 nm, 532 nm, 638 nm) and three single-photon-sensitive avalanche photodiodes with bandpass filters (APDs, APD1: 500-550 nm, APD2: 575-625 nm, APD3: 635-835 nm). For data acquisition, a 532 nm excitation laser was used at 10% power, along with the kymograph function of the software used to run the Lumicks C-trap, Bluelake. Fluorescence was detected using avalanche photodiodes APD2 and APD3. The kymograph settings included a pixel size of 100 nm, a pixel dwell time of 0.3 ms, and a line scan time of 11.9 ms. Analysis of force data was done as described before (Mondol et al., 2023), but with an expected DNA contour length of 1735 nm, due to the use of longer DNA handles in the current work.

## SUPPLEMENTARY MATERIAL

Amino acid sequence of the protein construct; procedure for the precise adjustment of the confocal plane; example of force readout for non-fluorescently labelled Hsp90 in the presence of AMP-PNP; kymograph recordings including the scanned area of the beads (PDF)

## Supporting information

Amino acid sequence, experimental procedures

## ACKNOWLEDGMENTS

KT is supported by an MSCA-IF (NOTE, grant agreement number 101028366) and by the NWO grant (Solve90, grant agreement number OCENW.M.22.092). Special thanks to Dr. Christopher Battle and Tycho Marinus for their invaluable assistance in providing and improving data analysis tools, to Dr. Marco Simonetta and Dr. Bärbel Lorenz for their advice and to Ronald van der Meulen and Prof. Giovanni Maglia for a critical reading of the manuscript and for their highly insightful comments and support.

