## Supplementary material for "One-Pot Dual Protein Labelling for Simultaneous Mechanical and Fluorescent Readouts in Optical Tweezers": Amino acid sequence, experimental procedures

### Supporting Information

#### Amino acid sequences of the protein used in optical tweezers experiments

The Hsp90 construct from *Saccharomyces cerevisiae*, also known as Hsp82, contains an amber codon mutation at position 452 (D452TAG) in the middle domain (shown as *, coloured red). This mutation allows for the incorporation of the non-canonical amino acid cyclopropene-L-lysine, which is used for labelling with DNA oligonucleotides. The N-terminal domain contains a cysteine at position 61 (D61C, coloured yellow) to enable labelling with fluorescent dyes. The construct also includes an additional C-terminal alpha helix (coloured purple), which has a strong tendency to form a coiled coil. This C-terminal alpha helix contains a cysteine (A6C, coloured yellow) that can form a disulfide bridge upon coiled coil formation, covalently linking the two Hsp90 monomers. Furthermore, the construct features a C-terminal hexa-histidine (His) tag (coloured green) for purification using a Ni-NTA column.

MASETFEFQAEITQLMSLIINTVYSNKEIFLRELISNASDALDKIRYKSLSDPKQLETEP
CLFIRITPKPEQKVLEIRDSGIGMTKAELINNLGTIAKSGTKAFMEALSAGADVSMIGQ
FGVGFYSLFLVADRVQVISKSNDDEQYIWESNAGGSFTVTLDEVNERIGRGTILRLFL
KDDQLEYLEEKRIKEVIKRHSEFVAYPIQLVVTKEVEKEVPIPEEEKKDEEKKDEEKK
DEDDKKPKLEEVDEEEEKKPKTKKVKEEVQEIEELNKTKPLWTRNPSDITQEEYNAF
YKSISNDWEDPLYVKHFSVEGQLEFRAILFIPKRAPFDLFESKKKKNNIKLYVRRVFIT

DEAEDLIPEWLSFVKGVVDSEDLPLNLSREMLQQNKIMKVIRKNIVKKLIEAFNEIAED

SEQFEKFYSAFSKNIKLGVHEDTQNRAALAKLLRYNSTKSV*ELTSLTDYVTRMPEH

QKNIYYITGESLKAVEKSPFLDALKAKNFEVLFLTDPIDEYAFTQLKEFEGKTLVDITK

DFELEETDEEKAEREKEIKEYEPLTKALKEILGDQVEKVVVSYKLLDAPAAIRTGQFG

WSANMERIMKAQALRDSSMSSYMSSKKTFEISPKSPIIKELKKRVDEGGAQDKTVK

DLTKLLYETALLTSGFSLDEPTSFASRINRLISLGLNIDEDEETETAPEASTAAPVEEV

PADTEMEEVDPGEQKCEEWKRRYEKEKEKNARLKGKVEKLEIELARWRPGSAWS

HHHHHH

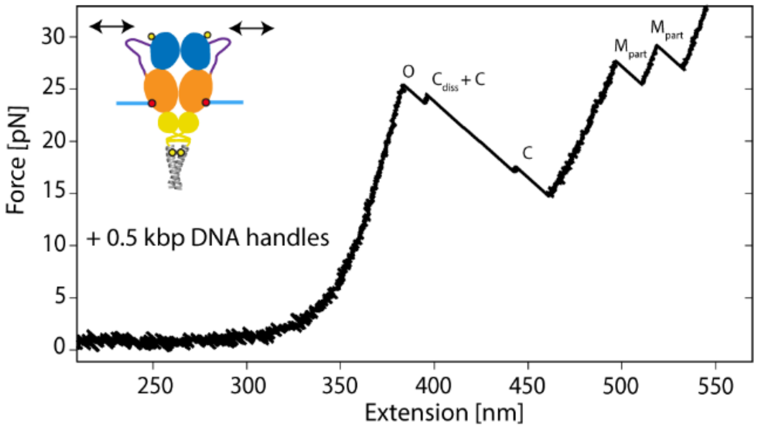


**Supporting Figure 1**: **Force readout for non-fluorescently labelled Hsp90 in the presence of AMP-PNP**. Example of a constant velocity unfolding trace at 500 nm/s for non-fluorescently labelled Hsp90 with shorter 0.5 kbp DNA handles, as used in Tych et al., (2018), in the presence of AMP-PNP. The trace depicts a change in length corresponding to the transition from the closed N-terminal conformation to the open state (O), followed by the dissociation of the Hsp90 C-terminal domains ($C_{diss}$), the unfolding of both C-terminal domains (C), and ultimately the unfolding of both middle domains ($M_{part}$).

#### Confocal plane adjustment and detection of fluorescence

To allow the detection of fluorescent dyes attached to Hsp90 using a hybrid C-Trap instrument, precise alignment of the confocal focus (controlled by the objective position) with the optical trap focal plane (where the beads, with the tethered sample, are trapped) is essential (Figure S2A). To achieve this alignment, SYBR Safe stain was employed to visualize the DNA handles that tether the dye-labelled Hsp90 protein between the beads (Figure S2B). This approach mitigates challenges associated with dye photobleaching that occur when relying solely on the fluorescently labelled Hsp90 constructs for alignment.

After aligning the confocal focus with the trap focal plane, the fluorescent dyes Atto-550 and Atto-647N, attached to Hsp90, were visualized upon excitation at 532 nm in presence of AMP-PNP (Figure S2C). It should be noted that the labelled construct with attached DNA handles was pre-incubated with anti-digoxigenin polystyrene beads. Therefore, fluorescence is visible not only from the construct tethered between the two beads but also from constructs attached to the anti-digoxigenin bead.

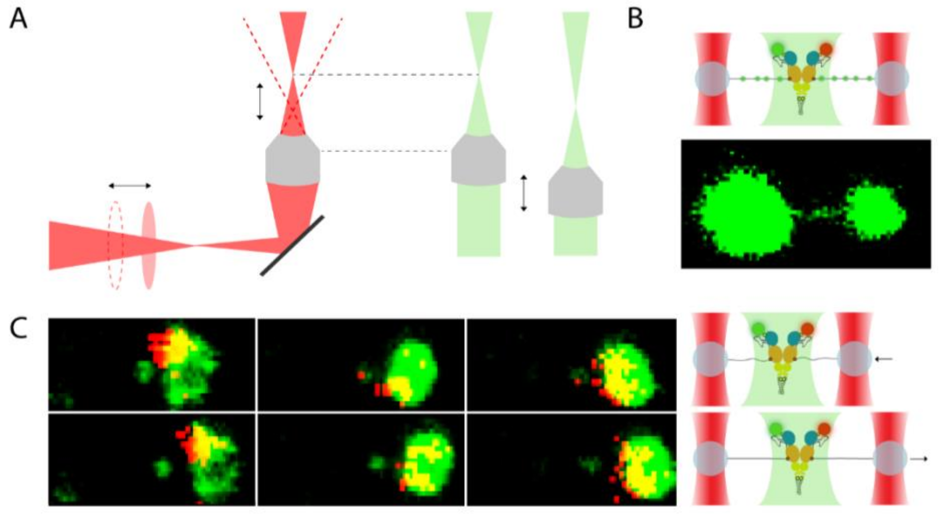


**Supporting Figure 2: Adjustment of the confocal plane and fluorescence detection.** (A) Illustration showing the alignment of the confocal focus (green) determined by objective positioning, with the optical tweezers trap focal plane (red). (B) SYBR Safe was used to visualize the DNA handles tethering the dye-labelled Hsp90 construct, aiding in the alignment of the confocal focus with the trap focal plane. (C) Fluorescence signals in presence of AMP-PNP were detected from Atto-550 and Atto-647N dyes attached to Hsp90, which is tethered between two beads. Signals were also detected from Hsp90 constructs attached to the anti-digoxigenin bead on the right side in each displayed image scan. The instrument used has an offset between the green and red channels. This offset was not corrected in the image scans shown here but was later addressed in the kymographs using a custom Python script to determine the pixel shift between the two channels.

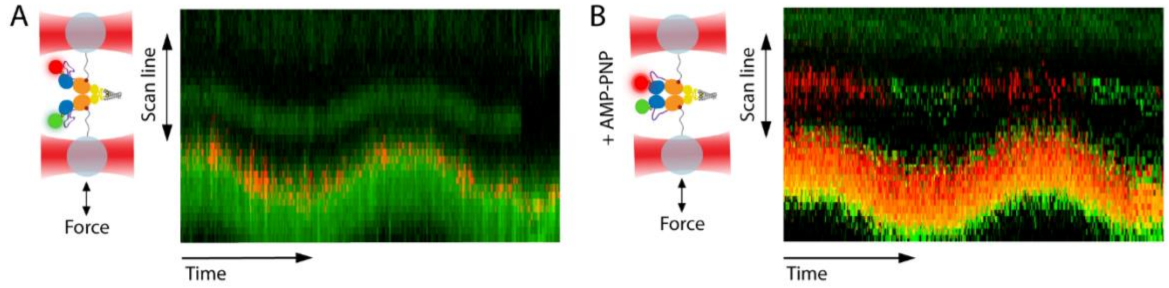


**Supporting Figure 3**: **Kymograph recordings**. Kymographs recorded in the absence (A) and presence (B) of AMP-PNP, including fluorescent signals from the polystyrene beads used in the experiment. These beads exhibit green autofluorescence when excited at 532 nm. The labelled construct, with attached DNA handles, was pre-incubated with anti-digoxigenin polystyrene beads, which are visible at the bottom of the kymographs. Consequently, red fluorescence is observed not only from the construct tethered between the two beads but also from constructs attached to the anti-digoxigenin bead. The C-Trap instrument used exhibited a misalignment between the green and red channels. To correct this in the kymographs, a custom Python script was employed to calculate the pixel displacement between the channels. This adjustment was applied using Photoshop, which was also utilized to enhance brightness for improved visibility.
